# Biomolecular solution X-ray scattering at n^2^STAR Beamline

**DOI:** 10.1101/2022.05.20.492803

**Authors:** Oktay Göcenler, Cansu M. Yenici, Kerem Kahraman, Cengizhan Büyükdağ, Çağdaş Dağ

## Abstract

Small angle X-ray Scattering (SAXS) is a method for determining basic structural characteristics such as size, shape, and surface of particles. SAXS can generate low resolution models of biomolecules faster than any other conventional structural biology tools. SAXS data is mostly collected in synchrotron facilities to obtain the best scattering data possible however home source SAXS devices can also generate valuable data when optimized properly. Here, we examined sample data collection and optimization at home source SAXS beamline in terms of concentration, purity, and the duration of data acquisition. We validated that high concentration, monodisperse and ultra pure protein samples obtained by size exclusion chromatography are necessary for generating viable SAXS data using home source beamline. Longer data collection time does not always generate higher resolutions but at least one hour is required for generating a feasible model from SAXS data. Furthermore, with small optimizations both during data collection and later data analysis SAXS can determine properties such as oligomerization, molecular mass, and overall shape of particles in solution under physiological conditions.

## Introduction

X-ray scattering is a widely used technique for characterizing chemical and structural properties of a variety of biological macromolecules and complexes. The characterizations of chemical composition, structure, and size can be done by utilizing X-rays and its scattering on a detector when X-ray beams hit electrons of an atomic shell (Kane et al., 1986). If the wavelength and the energy of the X-rays do not change but angle of X-rays change when X-rays are scattered from electrons, the scattering is classified as elastic scattering. In contrast, when energy of X-rays change after being scattered from electrons, the scattering is classified as inelastic scattering (Kane, 1992). Two elastic scattering methods, small angle X-ray scattering (SAXS) and wide angle X-ray scattering (WAXS), are the most commonly used methods for analytical characterization of biological molecules (Chavas, 2014). SAXS can be used to generate nanoscale resolution characterization of samples, whereas WAXS is used for atomic scale resolution characterization of smaller macromolecules as it covers wider angles (Makowski, 2010). The measurement of small angles for SAXS measurement is achieved by moving the detector away from the sample and the measurement of certain wide angles is achieved by moving the detector closer to the sample. In this article, usage of X-ray scattering method for biological molecule characterization in solution will be explained with exemplary data generated from a newly constructed small/wide angle X-ray scattering n^2^STAR beamline at Koç University.

As described above, small angle X-ray scattering (SAXS) is an established X-ray imaging method used in nanotechnology, chemical engineering, and structural biology as a tool for analyzing structural components of relatively larger molecules by their size, shapes, and morphology (Das and Doniach, 2008). More specifically in structural biology it is used to determine size, shape and the morphology of proteins and other biological macromolecules but combined with NMR it can generate 3-D protein structures. Despite providing very low resolution, SAXS is becoming an important method among structural biologists due to its swiftness and practicality. SAXS is also a reputable X-ray imaging method used for structural characterization of particles ranging in between 1 nm to 1000 nm (Pauw, 2013). As X-rays are scattered by Thomson scattering model, electrons are initially scattered elastically therefore resulting in larger molecules being scattered in smaller angles and making SAXS a competent method for analyzing especially larger molecules (Skou et al., 2014). Conceptually SAXS has similar characteristics with other X-ray imaging methods such as X-ray crystallography (XRC) but differs in the condition of the sample.X-ray Crystallography (XRC) exclusively requires particles ordered in a crystal lattice, in contrast, SAXS does not require such specific formation and SAXS samples are analyzed while target molecules are in solution (Grant et al., 2011). This characteristic of SAXS defines the primary difference between SAXS and XRC in terms of data analysis and strategies on the refinement of the data. In XRC, the diffraction data collected from crystallized samples appear in a discreet manner due to highly ordered localization of the molecules in the sample and generating higher and noticeable signals. Scatterings collected from XRC experiments can generate a high resolution 2-D image of a molecule from a certain angle and thus require multiple rotations to have a total 3-D understanding of the sample (Matthews, 1976). Although XRC images come with comparably higher resolutions, the structure obtained from XRC could be misleading since it requires fixing biological molecules with non-biologically relevant interactions under harsh conditions inducing hyperosmolarity of the environment. In contrast, diffraction data obtained by SAXS comes from randomly oriented and biologically free molecules or proteins which, unlike crystals, are not induced into static conditions. Flexibility and accessibility of SAXS as a method provide fast structural analysis of protein structures and certain characteristics of biomolecules such as formation of oligomeric protein structures and protein ligand interactions. Combination of SAXS and NMR spectroscopy for structural studies utilizing each method’s own advantages will result in generation of more accurate models.

Due to advances in small angle scattering (SAS) data acquisition and a variety of computational methods for analysis of this data, the volume of the papers published involving SAS data increased significantly. Consequently, a central data bank was required to store the data published to increase the availability of the results of these experiments to interested researchers. In response, The Small Angle Scattering Biological Data Bank (SASBDB) was established to meet the growing need to store ever-increasing SAXS/SANS data (Valentini et al., 2015). As a result of growing popularity of SAS data among researchers, it was decided to develop a universal data format to prevent spending time writing subroutines to utilize data in a variety of formats. Lacking a universal format of storing SAS data was a problem from the beginning of usage of SAS but it wasn’t until 1999 when a decision was taken to develop a universal format of SAS to make it more available to newer researchers in this growing field (Malfois and Svergun, 2000). In response to this problem an extension to Crystallographic Information File (CIF) developed which is called small angle scattering Crystallographic Information File (sasCIF) for storing one-dimensional SAS data along with certain parameters of the experiment (Kachala et al., 2016).

After SAXS’ initial introduction to structural biology as a tool, a variety of new methods have been developed for SAXS to analyze structural properties of biological macromolecules in solution and under physiological conditions. Among one of the useful methods for analysis of protein oligomeric structures is the prominent application of SAXS, since in solution properties of proteins differ for individual proteins but could be amplified in presence of multiple proteins expected to form oligomeric structures particularly in solution (Korasick and Tanner, 2018). As concentrations of each protein in a complex can be adjusted, unlike conventional co-crystallization methods, SAXS allows greater control in important parameters allowing determination of a dynamic self-association equilibrium. As previously mentioned, SAXS data is often combined with higher resolution structural biology tools to obtain sufficient details to generate a 3D model of a protein or a complex. However, a computational protocol called ATTRACT-SAXS was developed which can generate high resolution structures of protein-protein complexes by itself. (Schindler et al., 2016) Intrinsically disordered proteins (IDP) and intrinsically disordered regions of proteins have also been a problem for X-ray crystallography as static models generated by crystallized protein diminish the possibility of structural characterization that are close to actual native conformations of proteins. In that sense, integrative studies with SAXS, NMR and single-molecule Förster Resonance Energy Transfer (smFRET) are capable of uncovering functions of intrinsically disordered regions of protein and more realistic and collective understanding of collection IDP conformations obtained by these methods (Gomes et al., 2020).

Crystallization of large biomolecules like antibodies for their structural characterization remains a challenge for structural biologists mainly for drug development studies. Understanding the structural characteristics of antibodies is a critical step for developing more competent therapeutic agents and SAXS could be utilized as a high throughput tool for skimming general structural characteristics of antibodies. In relation to singular characterization of antibodies, real time analysis of supplementation of additional compounds to determine the effects of these compounds or interacting particles paves the way for development of more efficient antibodies (Belviso et al., 2022). It is also shown that SAXS is capable of being a high throughput protein structure analysis tool with an automated system, without requiring extensive crystallization of multiple protein samples. Despite providing structures with atomic accuracy, X-ray crystallography has major limitations on determining structures of intrinsically disordered or flexible regions of proteins; however, SAXS provides valuable effort to structural genomic analysis in addition to conventional present tools in this field (Hura et al., 2009). SAXS perhaps on its own is not always sufficient for generating highly detailed structructures of large RNAs, can be combined with refined NMR structures to validate effectiveness of SAXS for assembling RNAs complexes observed in ribosomes. Structural models obtained from diffraction of ribosome crystals showed similarity between the existing model and thus effectiveness of using SAXS integrated with collection of high resolution NMR structures of RNAs for conformational sampling (Dagenais et al., 2021).

During early years of SAXS data collection, a variety of decentralized tools were developed for analysis of scattering data but ATSAS 3.0 software is the most widely used and the unarguably the most comprehensive software in terms of available programs for building models and visualization of small and scattering data. Due to growing availability of SAXS data in Synchrotron facilities, more analysis tools have been developed including: Scatter (Förster et al., 2010), BornAgain (Pospelov et al., 2020), SasView (Tan et al., 2021), and Irena (Ilavsky and Jemian, 2009). The available programs and tools in ATSAS are mainly developed for analysis of biological macromolecules and complexes in solution, however some programs are suitable for monodisperse and polydisperse systems in general (Manalastas-Cantos et al., 2021). In addition to model building models, ATSAS package also provides a handy GUI for manual data analysis called PRIMUS primarily utilized for basic data manipulation. Obtained SAS data can be directly loaded into PRIMUS and essential analysis performed by generation of Guinier and Porod plots to derive basic parameters such as radius of gyration (Konarev et al., 2003). Evaluation of the particle distance distribution function P(r) is critical for Ab initio modeling and GNOM is used as an indirect transform program using 1-D SAS data as input (Svergun, 1992). ATSAS offers a variety of programs for ab initio modeling for deriving low resolution three dimensional shapes from small angle X-Ray scattering of randomly oriented biological macromolecules in solution. DAMMIM, an ab initio modeling program in ATSAS package, uses a dummy atom model for building a multiphase model of a biological macromolecule. Initially a spherical space is filled with spherical dummy atoms in search volume, and the model is reconstructed according to the scattering data via characterizing each dummy atom solvent or scattering belonging to the molecule (Svergun, 1999). Additionally for rapid ab initio shape determination, DAMMIF is developed by utilization of DAMMIM program for accelerated model generation from small angle scattering data (Franke and Svergun, 2009). For molecules with experimentally determined atomic structures, CRYSOL is employed to fit small angle scattering curves of particles in solution into experimental coordinates of atoms obtained by X-ray crystallography (Svergun et al., 1995) and later structures obtained by NMR (Mertens and Svergun, 2017). Collective application of these programs provided by ATSAS is sufficient to build a three dimensional model from the original small angle scattering data comparable with higher resolution models obtained from X-ray crystallography or NMR.

Our state of the art research facility (n^2^STAR) is equipped with Anton Paar SAXSpoint 5.0 (Figure 1) with the Primux 100 high-performance 100 micro Cu microfocus sealed-tube X-ray source and 2D 1M EIGER2 R HPC detector andTrueSWAXS, allowing to choose the optimum q-range for your experiment over a large sample-to-detector distance (SDD) range from ≤45 mm to >1600 mm.

**Figure 1.**
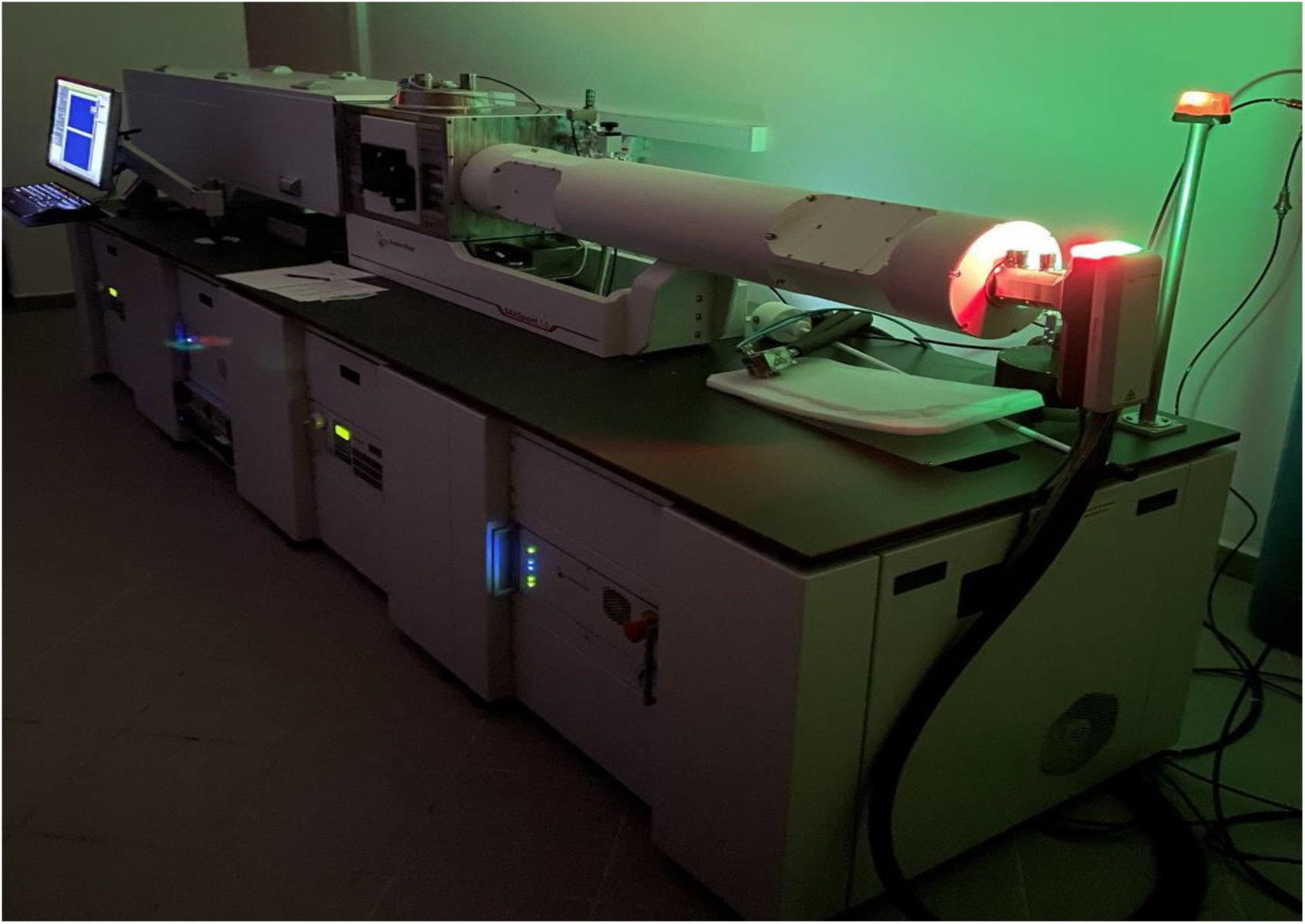
n^2^STAR SAXS/WAXS/GISAXS beamline.

## Material and Methods

### Protein expression and purification

Genes encoding recombinant proteins (UbcH8, UbcH7, E6AP HECT) were cloned into N-terminal GST-tag containing pGEX expression vectors for previous studies by our collaborators (Tokgöz et al., 2012; Ronchi et al., 2013). Following transformation into *E. coli* BL21(DE3) cell line, the cells were grown on antibiotic containing agar plates. 50mL and 1L Luria-Bertani (LB) medium were prepared and autoclaved. Selected colonies were transferred into 50 mL LB media containing the appropriate antibiotics and grown overnight in the temperature controlled shaker set to 37°C and 110 rpm. 50 mL LB cultures were transferred into large flasks containing 1L LB medium. Large scale cultures were grown in the incubator set to 37°C and 110 rpm. The incubation temperature was lowered to 16°C after OD at 595 nm exceeded 0.3 A. Protein production was induced by 0.4 mM (0.4 μM final concentration) IPTG at 0.8 OD. Cells were harvested by centrifugation at 3500 rpm for 45 minutes, using a swinging bucket rotor centrifuge. The cells were lysed using a sonicator after the addition of lysis buffer (50 mM Tris, 500 mM NaCl, 0.1% Triton X-100, 5% glycerol, 1 mM DTT, pH 7.5). Cell lysates were centrifuged at 20000 rpm for 1 hour and the supernatant was collected. GSTrap columns were equilibrated using the Tris-based buffer (50 mM Tris, 150 mM NaCl, 1 mM DTT, pH 7.4) and the supernatants were loaded at 1 ml/min. After washing the column bound proteins with the same Tris-based buffer, the proteins were eluted with the elution buffer (30 mM glutathione, 50 mM Tris, 150 mM NaCl, 1 mM DTT, pH 7.4). The proteins were diluted 1:2 using the wash buffer and the proper restriction enzyme was added for each protein to cleave the GST tags (Tokgöz et al., 2012; Ronchi et al., 2013). Glutathione was removed from the protein solution with overnight dialysis using 3 kDa cut-off dialysis membrane before reverse GSTrap. Samples were loaded to GSTrap columns equilibrated using the wash buffer. Free proteins were collected in the flowthrough and GST tags were eluted using the elution buffer. GST tags collected from the purifications were pooled together and concentrated using a 10kDa cutoff centrifugal filter. Size-exclusion chromatography (Superdex S200 Cytiva) was performed to isolate ultra pure GST protein. The column was equilibrated using (50 mM Tris, 150 mM NaCl, 1 mM DTT, pH 7.5) and the sample was injected to the column. Fractions containing ultra pure GST were determined by SDS-PAGE. The fractions were combined and concentrated to 4 mg/ml using 10 kDa cutoff centrifugal filters for SAXS measurements.

### SAXS beamline preparation

In order to start the device, first of all the main switch is turned on which is located on the back of the device. To gather data, the gap between the sample and the detector must be kept at a low vacuum to avoid air scattering. This air scattering can be apparent when it is compared with the low scattering signal of macromolecules. For this purpose, vacuum pumps are switched on (Figure 2A). The less amount of air results in the decrease of the direct interaction of the air with the direct beam coming from the light source. After the vacuum pumps and chillers (Figure 2B) are turned on, which are used to adjust the temperature, the X-ray source/the generator (Figure 2C), the detector controller (Figure 2D), computer and the sample unit controller (Figure 2E) are switched on respectively.

**Figure 2.**
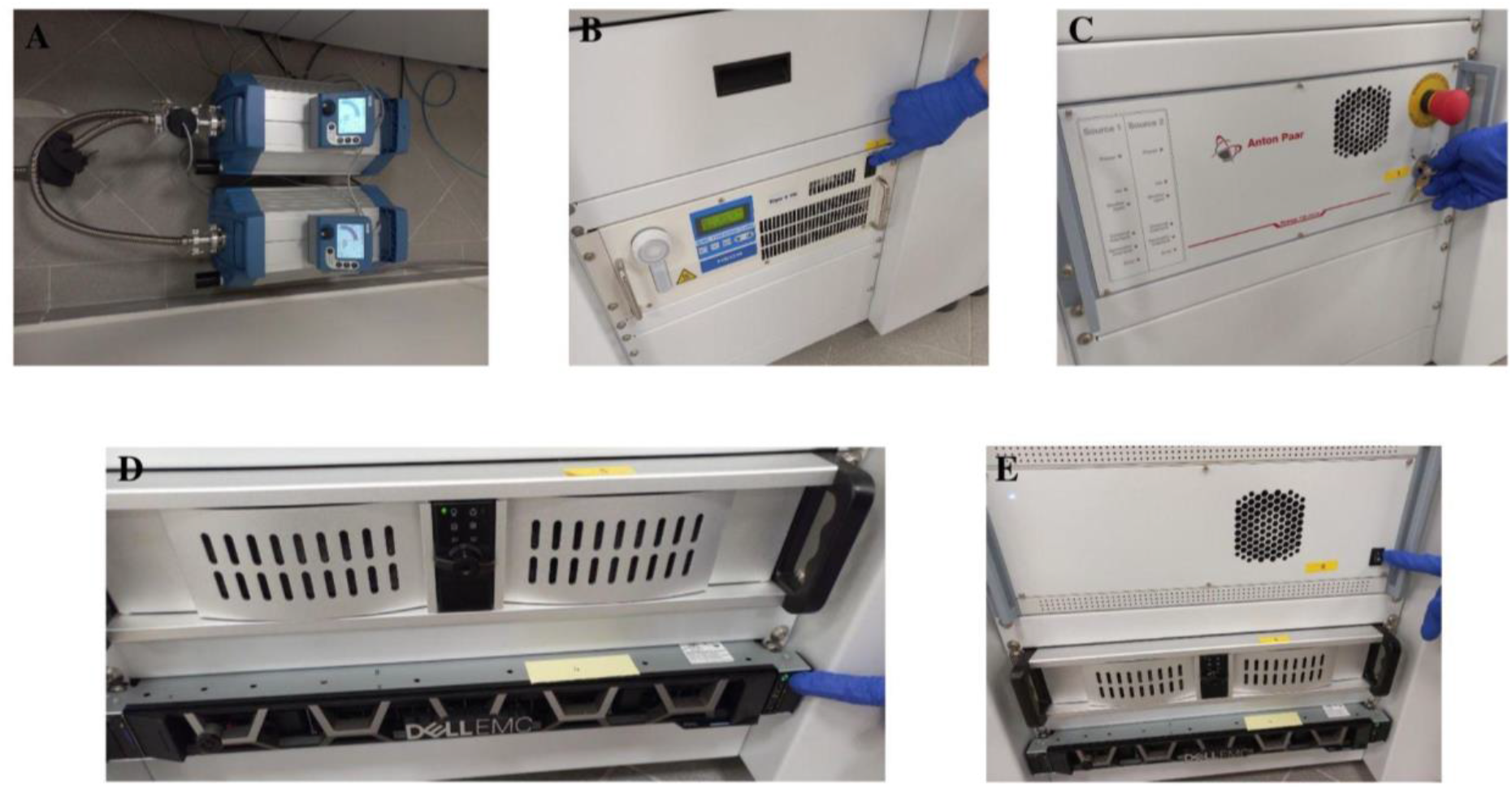
SAXS beamline preparation steps.

### Sample Load

There are two possible setups for biomolecular data collection in solution. First one is the static cell; when a static cell measurement is performed, the loaded sample is exposed to X-rays for some amount of time. This option may be preferred if there is a small amount of sample, the sample is obtained in a difficult way, the sample is expensive or it is scattering strongly. Second option is the flow cell. During the flow cell measurement, the sample that is exposed to X-ray beams is exchanged with a fresh sample. This measurement type may cost more or necessitate a larger amount of sample but it provides longer measurements without radiation damage and also it makes it possible to measure the signals that are weakly scattered. In our case, the sample is loaded to the static quartz tube (Figure 3A) using a micropipette. Before loading the sample, the capillary tube is washed with distilled water and dried using dry air for better results as there might be some deposits on the sample cell walls. After the washing, the sample is loaded (Figure 3B). It is important not to have air bubbles left inside the capillary tube. Once the sample is loaded, one side of the tube is closed with one finger while closing the other side of the tube. Then the other side of the tube is closed (Figure 3C). The quartz tube is then placed in the sample holder. Heated/Cooled sample holder (Figure 3D) is used to adjust the temperature. Data from sample and buffer (base line) should be collected using the same exposures, in the same cell and in the same position. Additionally, it is recommended to start with the buffer data acquisition and then continue with the biomolecular sample using the same data collection parameters. The system is evacuated after placing the tube into the sample holder in order to obtain vacuum conditions. When the required pressure value is reached, the system is initialized to align the system components within the device for data acquisition.

**Figure 3.**
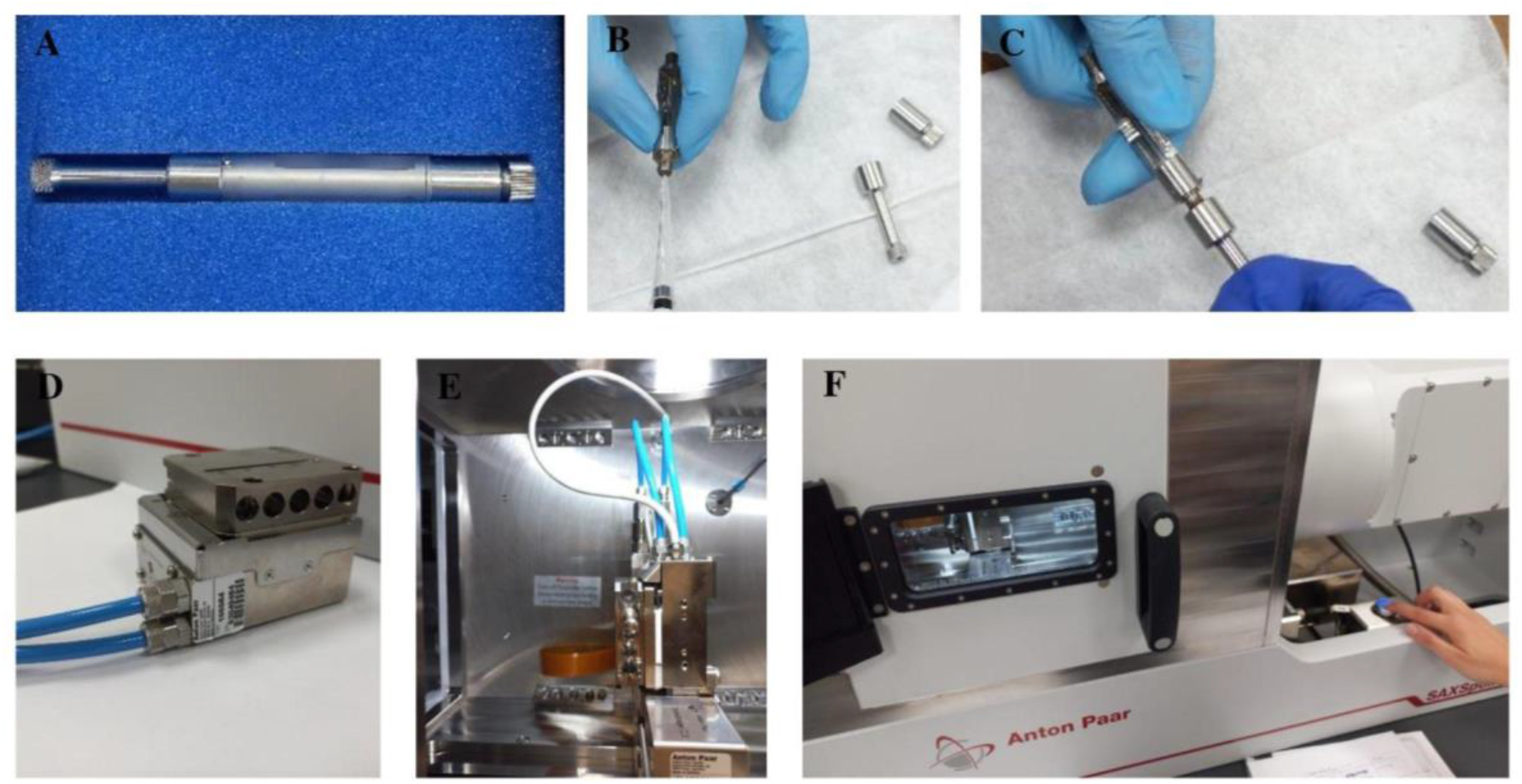
SAXS sample load.

### Data Acquisition and Processing

Before the acquisition, the temperature is adjusted to 10 °C. Sample/detector distance (SDD) is set to 1600 mm for SAXS experiments. Furthermore, the position of the beam stopper is checked and it is appropriately aligned with the incoming beam. The acquisition time and the number of frames are indicated. The transmission measurement is included in the data acquisition. After the data collection, SAXS output file (.dat) is obtained. This file needs to be processed in order to obtain the scattering pattern of the sample. For this purpose, the output file is uploaded to the SAXS analysis program and by clicking on “Source data”, the data files that are used are chosen. These files are the output file for the protein sample and the output file for the buffer/background. The first process is created and all data files are selected to be processed (Figure 4). First tool that is used is “Zero Point by Moments 2D” to select the primary beam coming from the X-ray source. Then “Masking” is performed in order to mask out the beam stop to exclude its influence on the sample data. To include transmission measurement into data acquisition, “Transmission” option is selected in the created “Q Transformation” tool. In “Data Reduction Step”, the whole data that is processed is chosen by entering the appropriate range values for x,y and angle parameters. The data is radially integrated. In the first “Standard Operations 1D” tool, transmittance normalization is done. This way the amount of X-rays that are passing through the sample is compared to when there is no sample inside by selecting the “Transmittance” option again. After these steps, another process is created to now process the obtained 1D data. In order to subtract the background scattering from the scattering of the sample in the buffer, “Standard Operations 1D” tool is created once again. The lastly processed background scattering data is selected as the “Reference background”. As the last step, the data file is obtained from the SAXSanalysis program to be compatible with the ATSAS program. Other file types that are compatible with other processing and analysis programs are available. The appropriate file type should be chosen according to the program that will be used for data processing.

**Figure 4.**
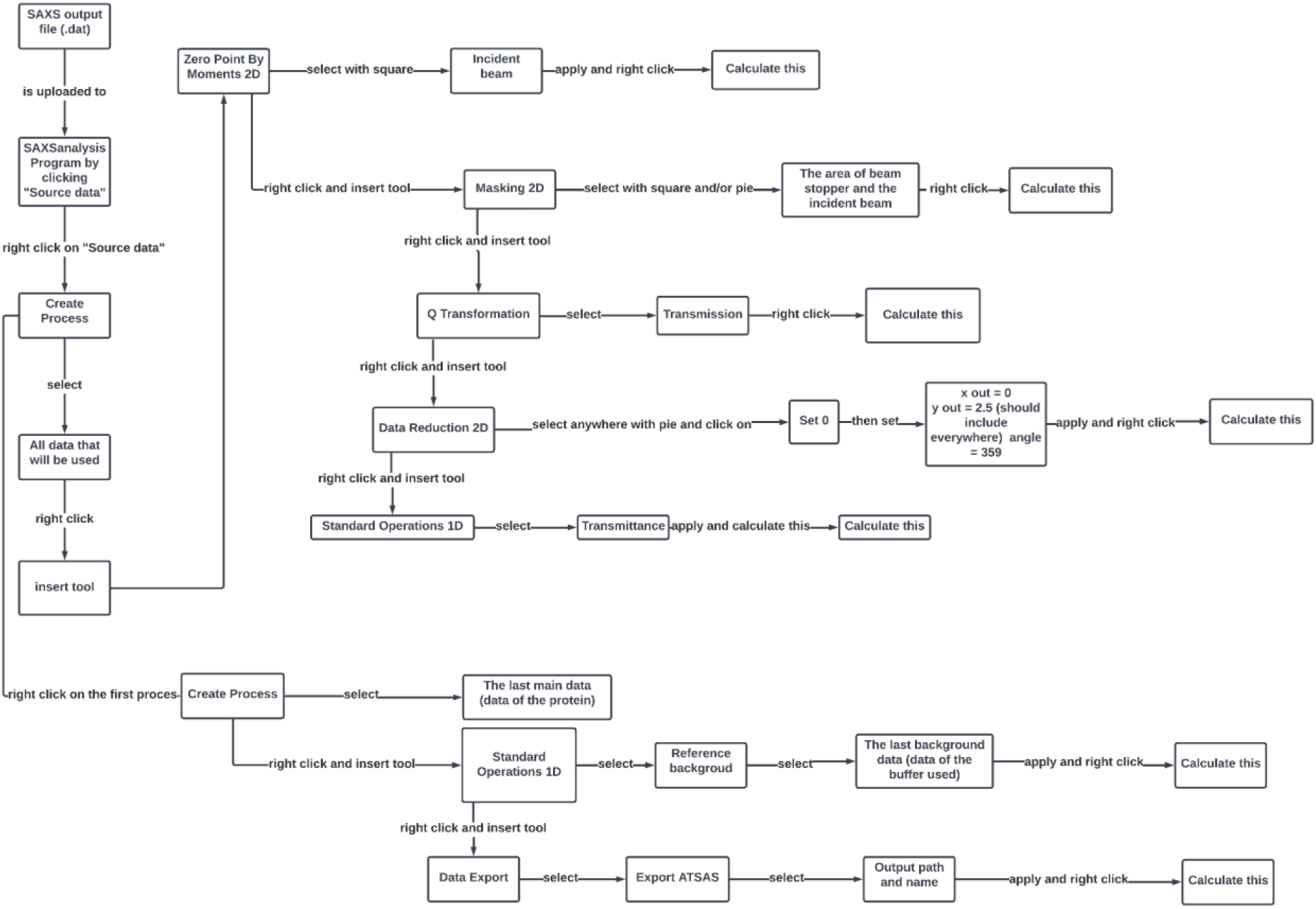
Data processing flowchart.

### Data Analysis

The scattering pattern of the protein in the buffer solution is determined by subtracting the scattering pattern of the buffer solution from that of the protein in the buffer solution as indicated in Figure 4. In this way, only the distribution belonging to the protein is obtained. The output file containing the scattering information of the protein is saved in a file type that can be analyzed using the ATSAS 3.0 program (Manalastas-Cantos et al., 2021). Logarithmic graphs are obtained by loading the output file into PRIMUS package of ATSAS 3.0. It gives information about whether the protein has UV damage or is precipitated. If there is any radiation damage or precipitation, repetition of the experiment is recommended after the optimization of experiment parameters. The radius of gyration is found using Guinier analysis (Guinier, 1939) and Fourier transformation which are included in the PRIMUS package (Konarev et al., 2003). GNOM is used to determine the particle radius (Dmax) (Svergun et al., 1992). Packages in PRIMUS can be used to estimate the molecular weight of the molecule of interest. Using the DAMMIF program, the low resolution structure of the molecule is created by “ab initio” modeling method and DAMAVER is used to obtain an averaged and optimized model of all models created by DAMMIF (Franke and Svergun, 2009). Moreover, the CRYSOL program (Franke et al., 2017) is used to compare and check the matching of the created models with the structural information that is contained in the protein data bank.

## Results and Discussion

GST protein was expressed in BL21 cells as tagged to various proteins to be used for its affinity to glutathione in affinity column chromatography assisted purification of the targeted proteins. As the GST tag is cleaved after purification, GST was selected due to the excess supply and its well characterized dimeric structure. GST tagged proteins are purified with 5 ml GST columns and the elution is treated with Thrombin to cleave the GST from the target protein. Cleaved target proteins are separated from GST by reverse GST column chromatography (Figure 5). Free GST obtained from a variety of samples were mixed altogether in order to increase the final protein concentration for SAXS analysis. Preliminary data showed that protein purity is very important for SAXS experiments (data not shown). In order to improve the data quality, gel filtration chromatography applied to samples. Purity and concentration of the samples are validated by SDS PAGE.

**Figure 5.**
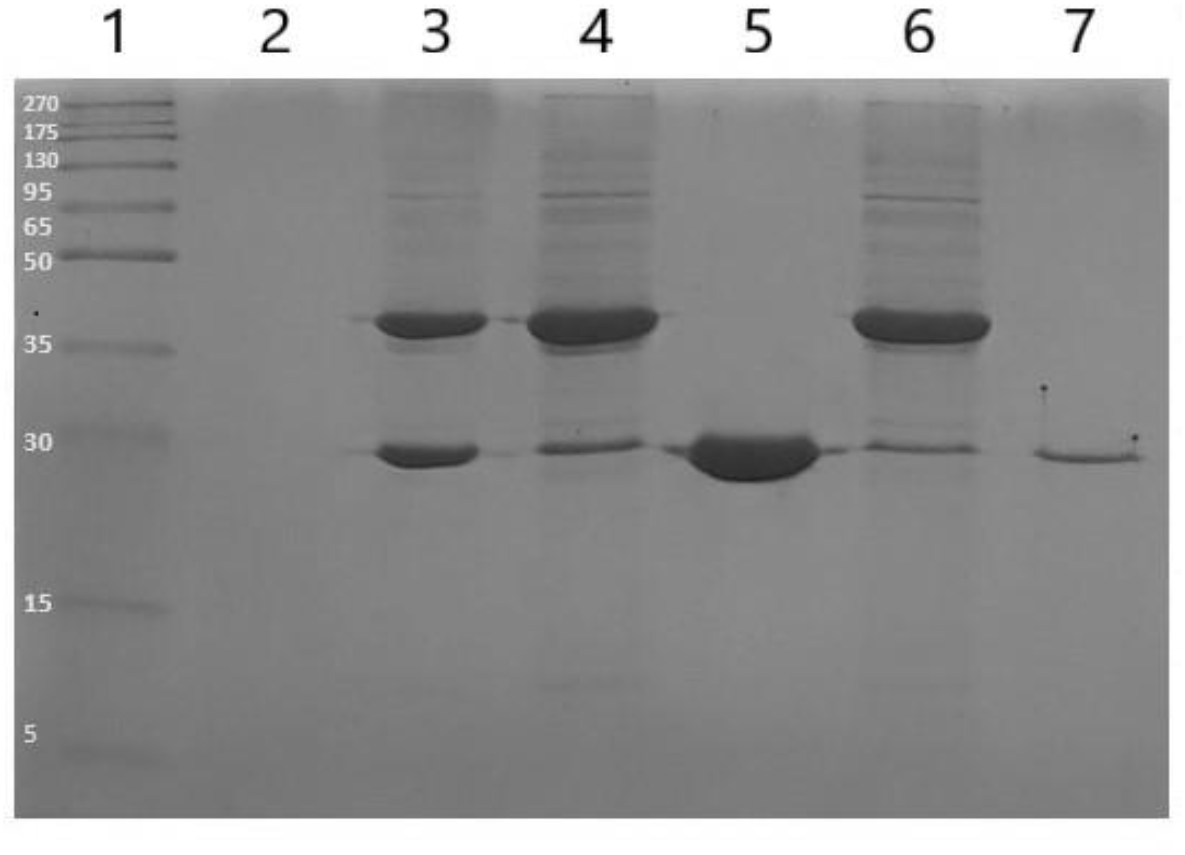
SDS-PAGE analysis of recombinant GST protein. (1) Marker (2)space before samples (3) E6AP_HECT after enzymatic cut (4) reverse GST flowthrough 1 (5) reverse GST elution 1 (6) reverse GST flowthrough 2 (7) reverse GST elution 2

Ultrapure free GST fractions from gel filtration chromatography were collected and concentrated to 4 mg/ml and 7 mg/ml levels with 10 kDa cutoff centrifugal filter. The optimal concentration for collecting small angle scattering data of free GST was not known and the effect of protein concentration to the quality of SAXS data was aimed to be determined by contrasting findings from both concentrations. According to the findings, there was a significant difference in the quality of the unprocessed SAXS data visibly observable on one dimensional data plot but also according to statistical indicators generated by PRIMUS data processing tool (Figure 6). The rest of the analysis was only performed with the data collected from the 7 mg/ml Free GST sample. Unlike inorganic compounds, biomolecules are highly sensitive to X-ray radiation generated by the light source; the quality of data collected is expected to decrease in a time-dependent manner. In order to check the effect of radiation damage to data collection, data was collected in 10 minute frames and before it was combined into one data file, possible alterations of the data was checked by superimposing data collected in each interval. During test runs with previous samples, one hour data collection was sufficient for generation of three dimensional models from the obtained data. However, collection time depends on a variety of factors ranging from concentration of the sample to aggregation or oligomerization of the proteins in solution. Nevertheless, one hour data collection works for the majority of proteins (data not shown). Radiation damage is not observed with any samples for one hour of data collection compared via 10 minute frames. Due to lack of data availability of protein SAXS data obtained from non-synchrotron based X-ray devices, there is no established protocol for data collection with weaker beamlines; however, data collection up to 5 hours with 30 minute intervals is possible without significant radiation damage in the sample (Živič et al., 2022).

**Figure 6.**
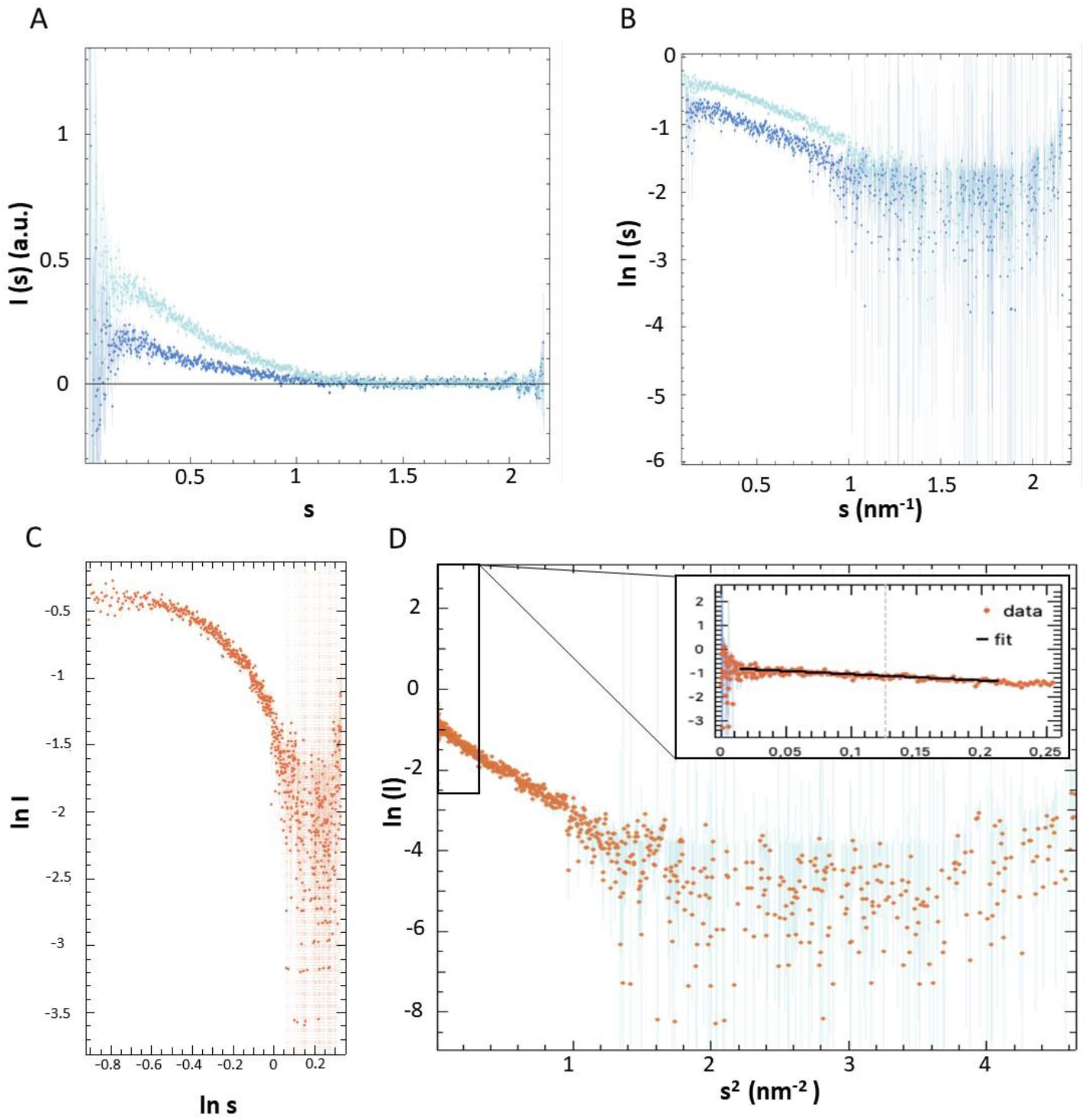
Solution SAXS analysis of the GST protein. (A) l(s) vs s plot of the experimental SAXS intensity obtained at 4.1 mg/ml (blue) and 7.0 mg/ml (turquoise) (B)ln I (s) vs s plot plot of the experimental SAXS intensity obtained at 4.1 mg/ml (blue) and 7.0 mg/ml (turquoise) (C) ln I vs ln s plot plot of the experimental SAXS intensity obtained at 7.0 mg/ml (D) Guinier plot obtained from 7.0 mg/ml sample

SAXS data obtained for the 7 mg/ml GST sample was analyzed using tools provided by PRIMUS in 3.0 ATSAS package. GST is naturally seen as a homodimer according to protein crystal structures thus the dimer form of GST was used for comparison with the model generated by SAXS. As SAXS data alone provides basic structural information according to the scattering profile with Guinier plots and Kratky plots (Figure 7), it was confirmed that GST was in fact in dimer form by comparing the estimated molecular weight from the data to the monomer of GST by PRIMUS. Radius of gyration is a valuable measurement calculated by the Guinier analysis as it specifies both weight and size of a particle around an axis without generating a model. In order to verify the dimer conformation of GST, Radius of gyration generated by PRIMUS (25.50) was compared to radius of gyration generated by CRYSOL analysis of dimeric form GST (22.48) (Table 1). Despite the slight difference between estimated and calculated radius of gyration values, the resemblance between Rg values is sufficient to confirm the small angle scattering is generated by a GST homodimer.

**Figure 7.**
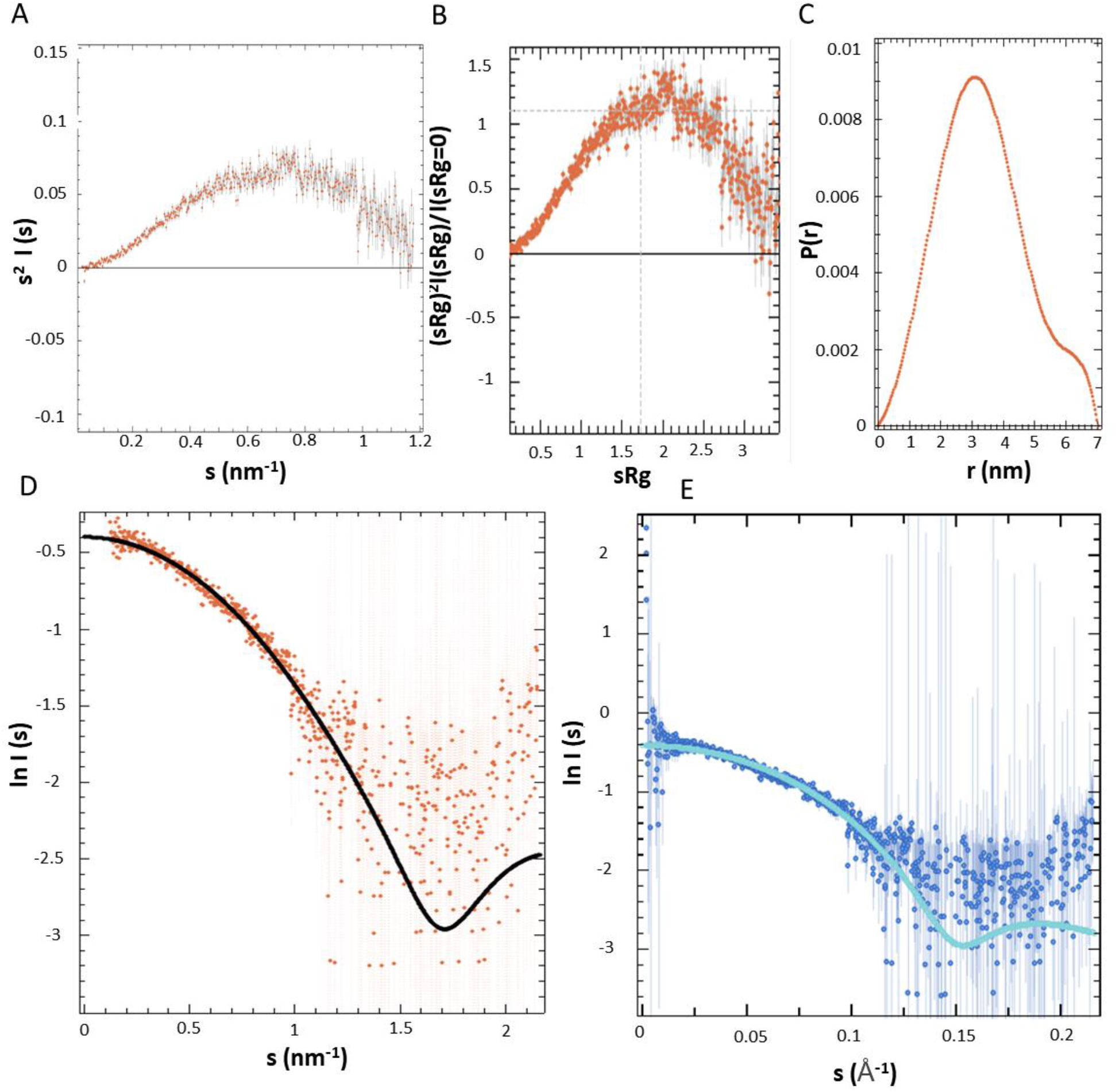
Kratky and CRYSOL analysis of the GST homodimer. **(A)** Kratky plot of free GST **(B)** Dimensionless Kratky plot of free GST **(C)** Distance distribution plot of free GST scattering data calculated by GNOM **(D)** the fit of the model to the experimental data calculated by GNOM **(E)** Fit of the calculated rigid body model by CRYSOL to experimental scattering (PDB: 1GNE)

**Table 1.** Molecular Size Parameters Obtained for GST from SAXS Data Analysis.

| Sample ID | R<sub>g</sub> (Å) | R<sub>g</sub><sub>cryst</sub> | V<sub>p</sub>, nm<sup>3</sup> | D<sub>max</sub>, nm | MMI(0) kDa | MM<sub>aa</sub> kDa | MM<sub>Porod</sub> kDa |
| --- | --- | --- | --- | --- | --- | --- | --- |
| GST | 25.50 | 22.48 | 80.20 | 7.03 | 59.30 | 55.80 | 58.80 |

Homodimer of free GST with the additional residues remaining from enzymatic cleavage of GST has a theoretical mass of 55.80 kDa. The molecular mass was estimated by ATSAS as 59.3 kDa and by Porod analysis as 58.8 kDa (Table 1). In spite of the difference in the mass estimation compared to the theoretical molecular mass, the error rates are not expected to exceed 10 % (Mylonas and Svergun, 2007) in this experiment. The molecular mass estimation error does not exceed 10 % in either estimation, also confirming monodisperse non-aggregated characteristics of our homodimer GST SAXS sample. Considering Radius of gyration and MM estimates were supporting a non-aggregated properly folded homodimer of GST molecules, Kratky plot was generated to investigate flexibility and disclose globular nature of the confirmed GST dimers. Folded and globular proteins expected have a Kratky plot with a well defined bell-curve that approaches zero after maxima at higher q values (Burger et al., 2016). In Figure 7A, the Kratky transformation is applied to GST SAXS data confirming folded and globular conformation of GST dimers as there is a clear bell curve with a peak around ∼2 sRg. In Figure 7B, dimensionless Kratky plot naturally gives the same result with a distinct bell curve that converges to 0 with increase of s. Pair distance distribution function P(r) is generated as another indicator of globular conformation of the scattered particle (Figure 7C)

In Figure 7E, a fitting curve is generated to validate the curve with high resolution models of GST dimers. As high resolution models of GST dimers exist, CRYSOL was run with a PDB containing a similar model with residual extensions due to cleavage from another protein (PDB:1GNE) (Figure 7E). According to CRYSOL analysis there are considerable similarities between the fitting curve of experimental data and the curve of GST homodimer generated by CRYSOL. As model-free analysis was satisfactory, a three dimensional model from experimental data was generated by using DAMMIF in the ATSAS package. DAMMIF was run 20 times and the average of these models was used to generate the most possible conformation of GST dimer in solution. Three dimensional model of GST homodimer (Figure 8) was aligned with the same model used for CRYSOL analysis. Remarkably, residual loop remaining after cleavage of GST is also detected by DAMMIF and generated as an extension to the globular model of the GST dimer (Figure 8). The residual loop on GST dimer is obtained from a static crystal lattice and the actual confirmation of this loop could be more flexible in solution, which SAXS is capable of detecting (Figure 8B).

**Figure 8.**
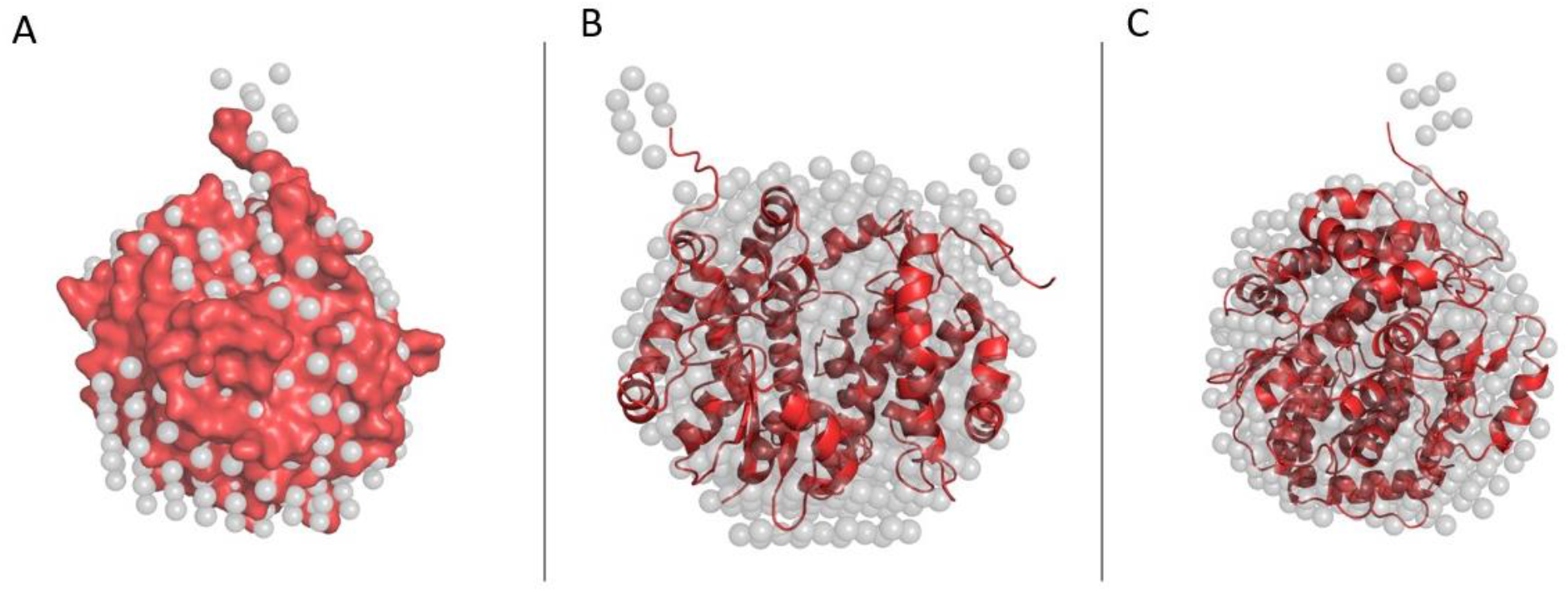
Reconstructed SAXS envelope of GST in dimeric form. (A) 3D reconstruction of dimeric GST superimposed with surface representation of high resolution PDB structure (B) a counterclockwise rotation of the protein around the vertical axis with 90-degree (C) 3D reconstruction of dimeric GST superimposed with cartoon representation of high resolution PDB structure

SAXS compared to conventional structural biology tools of X-ray crystallography, biological NMR spectroscopy is a tool capable of acquiring structural information about a biological macromolecule’s natural conditions in solution, without size limitation, and by matter of hours. Despite providing low resolution, SAXS can be a uniting ground for structural analysis of biological macromolecules as SAXS is capable of compensating the shortcomings of the other structural biology tools capable of generating structures with atomic accuracy. X-ray crystallography among structural biology tools is commonly used for generating protein models close to atomic accuracy. In order to obtain proper diffraction from biological molecules, proteins are fixed and ordered in a crystal lattice limiting flexibility and due to changing environmental conditions affecting certain conformational characteristics of protein when they are in physiological conditions (Putman et al., 2007). The data collection with SAXS only took an hour compared to necessary time for crystallization of the protein which can take months to achieve. SAXS’ role in structural biology would not be replacing X-ray crystallography but change how high throughput structural biology research takes place even in research centers without access to tools necessary for high throughput X-ray crystallography.

In a matter of hours, the dimeric form of GST was validated by a simple SAXS analysis, demonstrating effectiveness of SAXS for determining basic structural characteristics of biomolecules. Considering SAXS data was collected with a home-source X-ray, the data generated by this system is still capable of detecting and differentiating subdomains of fusion proteins in addition to oligomerization of proteins (Figure 9 A-N). Atomic resolution structures are certainly useful for a variety of studies in silico, however SAXS models alone could be sufficient to determine most basic characteristics of proteins if there is no obtaining high resolution structures. Furthermore integration of the conventional methods in structural analysis with SAXS could generate a considerable amount of data in a significantly short amount of time which could be useful as a high throughput analysis in the growing omics field of molecular biology. In recent years SAXS has been utilized for interdisciplinary structural biology studies and particularly useful as a quick analysis of large biological molecules of the stalk domain of SARS-CoV-2 spike protein (Živič et al., 2022). The environmental conditions including ligand or cofactor concentrations in solutions, temperature and pH can be manipulated in SAXS samples allowing detection of changes in the conformation of proteins in structural basis. Additional SAXS data including possible environmental changes is a good way of supporting the static X-ray crystal or cryo EM data to generate more reliable results in structural analysis research.

**Figure 9.**
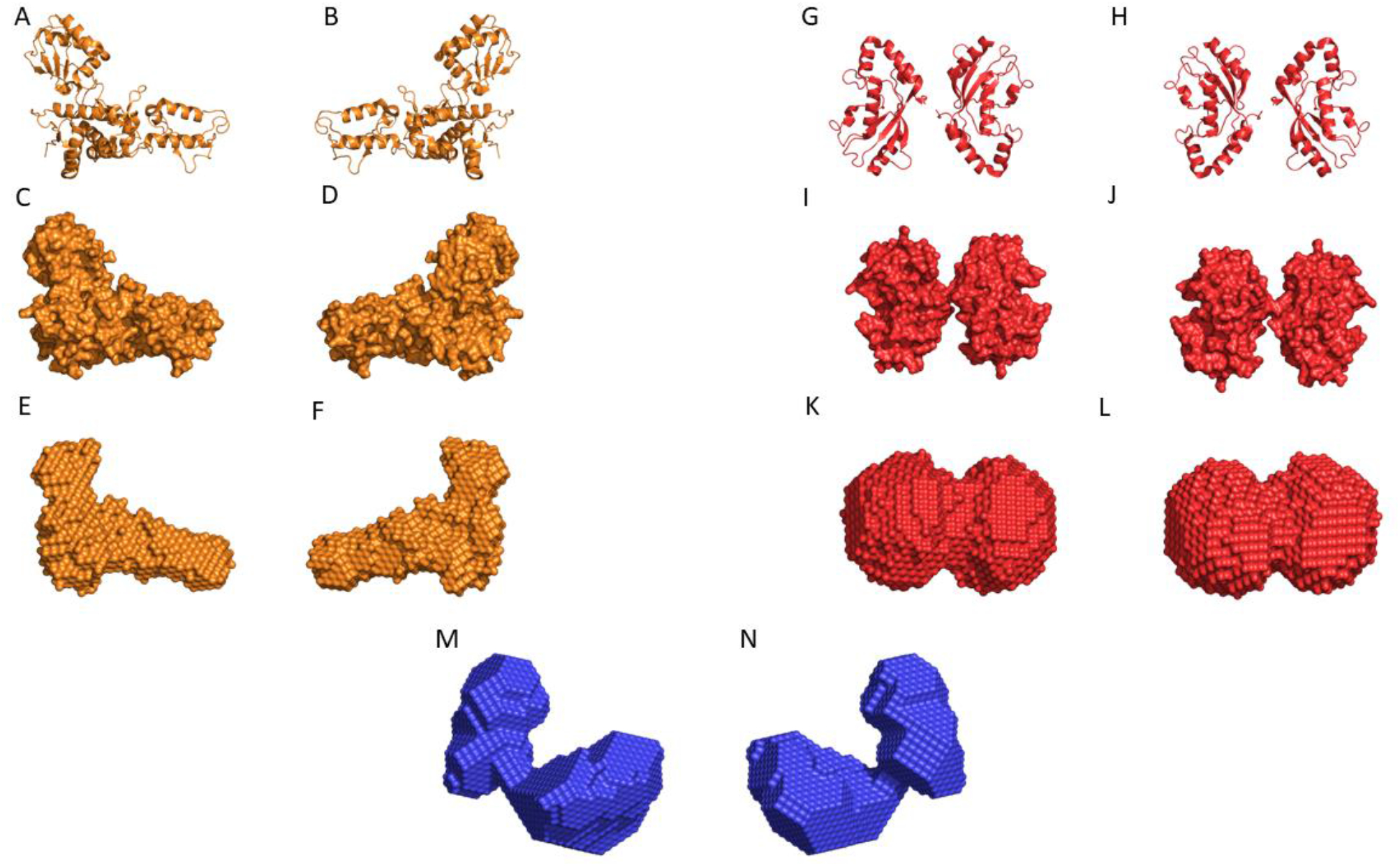
Examples of 3D reconstructions obtained from the data collected at n2STAR beamline. (A) and (B) Cartoon representation of target protein 1 PDB structure; (C) and (D)Surface representation of target protein 1 PDB structure; (E) and (F) Surface representation of target protein 1 (multi subdomain) model reconstruction from SAXS data (G) and (H) Cartoon representation of target protein 2 PDB structure; (I) and (J) Surface representation of target protein 2 PDB structure; (K) and (L) Surface representation of target protein 2 (dimeric) model reconstruction from SAXS data; (M) and (N) Surface representation of target protein 3 (fusion) model reconstruction from SAXS data

### Conclusion

SAXS is still not plug and play, and many parameters need to be optimized before getting successful results with particularly home-source SAXS devices. SAXS gives better results when the sample is ultra-pure, stable and the particles are in the same conformation. The set up for SAXS data collection may enhanced by including in line size exclusion chromatography to improve sample purity as a standard approach for home source SAXS data collection (Graewert et al., 2020). In addition to current data analysis and model construction algorithms, the growing field of deep learning is also contributing to the development of novel methods for more accurate three-dimensional model construction (He et al., 2020). Despite no deep learning approach being used in this study, it is likely as SAXS becomes more popular and deep learning algorithms developed for scientific data analysis, the approach of SAXS analysis of data can utilize such algorithms. Data entries to SAXS data bases with home source SAXS devices is still low, considering their potential benefits and its availability compared to a synchrotron facility. This number will increase day by day with the development of new generation detector technologies and new generation X-ray sources and better algorithms for analysis of SAXS data.

